# The dynamic climate adaptation of indicator warbler species complex in the Pacific Rainforest

**DOI:** 10.1101/2024.05.17.594794

**Authors:** Silu Wang

**Affiliations:** Department of Biological Sciences, University at Buffalo, 211 Mary Talbert Way, Buffalo, NY, 14260

**Author notes:** **Data management plan:** All the data involved in this study will be uploaded to Dryad.

**Keywords:** Climate change, Climate adaptation, plasticity, wood warbler, hybridization, evolution, temperate rainforest, *Setophaga*, niche competition

## Abstract

Understanding the functional responses of ecosystems to climate change is critical to predict and conserve global biodiversity. The evolution of indicator species provides an efficient avenue towards the functional biome response to climate change. Leveraging decades of citizen science data coupled with high-resolution spatiotemporal climate data, I dissected the climate change response of hybridizing *Setophaga* warblers, indicator species breeding in the mysterious canopy of the Pacific Rainforest. Breeding habitat temperature and precipitation powerfully predicted the breeding occupancy of the northern (*S. townsendi*) and the southern species (*S. occidentalis*). Both species showed positive climatic responses in the recent decade when the recent breeding occupancy was greater than the expected contingency. This implies the potential climate adaptation or life history plasticity in the breeding warbler populations. However, the *S. occidentalis* showed a compromised climate response in 2000-10, when the predicted breeding occupancy was significantly lower than the expected contingency. The compromised response might be due to rampant decline of July precipitation in 2000-10, compared to previous decades. Although the July precipitation continued to decline in 2010-20, *S. occidentalis* showed signs of recovery, after generations of climate adaptation. I further evaluated their breeding niche competition, reflected by the overlap of breeding occupancy between species. I found that the competition potential was the lowest in 2000-10 when both species were at the trough of breeding occupancy, which recovered in 2010-20. This study shed light on eco-evo feedback in the indicating species complex and illuminated the functional response of the rainforest ecosystem to climate change.

## Introduction

In the catastrophic climatic change (Kemp et al. 2022) and biodiversity loss (Brook et al. 2003; Dirzo and Raven 2003), efficient monitoring of the potential and trend in organismal response is crucial for conservation and restoration practice (Mawdsley et al. 2009; Dawson et al. 2011; Watson et al. 2012). The accruing citizen science data of organism occurrence (Johnston et al. 2023) and high-resolution spatiotemporal climate data collectively empower us to uncover organismal response to climate change.

The indicator species of vulnerable ecosystems are crucial stepping-stones for understanding the functional response of the ecosystems to climate change (Pearman et al. 2011; Siddig et al. 2016; Bal et al. 2018; Terrigeol et al. 2022). Here I leverage the indicating power of the wood warblers that breed in the canopy of the temperate rainforest of western North America as an avenue toward understanding the rainforest’s functional response to climate change.

The sister species, *Setophaga occidentalis* (SOCC) and *S. townsendi* (STOW) are indicator species of the old-growth rainforest (Carpenter et al. 2014) in western North America (Matsuoka et al. 1997; Huang 2013; Bielski et al. 2024). The temperate rainforest of Western North America has been experiencing unprecedented ecological vulnerability related to climate change (Coops and Waring 2011; Hicke et al. 2013; Scheller et al. 2018; Buotte et al. 2019; Goss et al. 2020). The breeding warblers hold nesting territories in the canopy of coniferous trees, such as Douglas Fir, Western Hemlocks, and Western Red Cedars (Barlow 1899), and their populations reflect the functional ecology of the temperate rainforest in Western North America (Bielski et al. 2024).

SOCC and STOW breed in divergent climatic conditions (Wang et al. 2021) yet hybridize and compete extensively at breeding overlaps (Rohwer and Wood 1998; Pearson 2000; Pearson and Manuwal 2000; Wang et al. 2019, 2020). The local mitonuclear ancestries of SOCC and STOW have been shown to covary with climatic variation in their breeding habitats (Wang et al. 2021). Whether SOCC and STOW are adaptively responding to the rapidly changing climatic conditions over the past decades remains an open question. Further, whether species-specific climatic response affects their breeding niche competition remains unknown, thus hindering the understanding of eco-evo feedback in rainforest climate change response.

I investigated the breeding warbler responses to climate change by comparing their expected breeding occupancy based on historical contingency and present breeding occupancy. If the warblers are positively responding to climate change, the present breeding occupancy should be greater than the expected breeding occupancy based on historical contingency. Evaluating the wood warbler’s response to breeding habitat climate change would shed light on the potential of climate adaptation of the temperate rainforest ecosystem. Further, I evaluated the effect of climate change on species boundaries as SOCC and STOW hybridize and compete upon contact (Rohwer and Wood 1998; Pearson and Manuwal 2000; Wang et al. 2019, 2020; de Zwaan et al. 2022). The geographic overlap in the predicted breeding occupancy of SOCC and STOW reflects the potential niche competition between the hybridizing species. A greater overlap suggests a stronger potential for competition in their breeding niches. Collectively, this time-series modeling synthesis sheds light on organismal, species boundaries, and ecosystem responses to climate change.

## Methods

### Breeding Site Climate

I downloaded the climate data from three decades: 1990-2000, 2000-2010, and 2010-2020 for western North America from WorldClim (Fick and Hijmans 2017; Harris et al. 2020). To represent the total climatic conditions at the breeding sites, I incorporated all the monthly climate data from CRU-TS-4.06 by using WorldClim 2.1 for bias correction with a spatial resolution of 2.5 minutes (∼21 km^2^ at the equator). At this temporal and spatial resolution, the climate variables available were: (1) average minimum temperature (°C), (2) average maximum temperature (°C), and (3) total precipitation (mm).

### Warbler breeding occurrence

I downloaded all the historical occurrences for *Setophaga occidentalis* and *S. townsendi* from GBIF.org (07 January 2024, GBIF.Org User 2024a,b). To focus on breeding individuals, I extracted occurrences for June each year between 1990 to 2020. To alleviate any identification error from the inexperienced observers, I excluded data from observers with less than 3 submission records.

### Breeding occupancy model

To train the breeding occupancy model for each decade (i.e. m_[1990-2000],_ m_[2000-2010],_ m_[2010-2020]_), I used the breeding season climate data to predict species presence/absence for each decade. The presence data was taken from the filtered breeding occurrence data from GBIF (see above). For absence data, I randomly generated pseudo-absence points (of equal sample size to the presence data) within the square of -155° to -105° W and 30° to 65° N. The collection presence/absence data was randomly divided into 5 folds so that ⅘ of it was used to train the occupancy model and the remainder ⅕of the data was saved as testing data. With the training data, I fitted the binomial generalized linear model in which the presence/absence is predicted by the interactions of all three climate variables. With the testing data for each model, I calculated the corresponding threshold probability of breeding presence with *pa_evaluate* function in the *predicted* package in R (Fielding and Bell 1997; Liu et al. 2011). I evaluated the model prediction accuracy as the following: (true positive + true negative)/total testing size.

### Breeding warbler response to climate change

To evaluate whether the wood warblers are responding to climate change, I calculated the *expected breeding continency* of the future decade (e.g. 2000-2010 or 2010-2020) with a historical occupancy model (m_[1990-2000]_) and climate input from the respective future decade (**Fig. 1**). At the same time, I calculate the *predicted breeding occupancy* with the species distribution models fitted with the climatic variation and species occurrence of each future decade. I compared the *expected breeding contingency* (*Exp*) and *predicted breeding occupancy* (*Pred*) within 2000-10 or 2010-20. A smaller value in the former than in the latter could suggest a positive response to climate change (e.g. signature of climate adaptation).

**Fig. 1.**
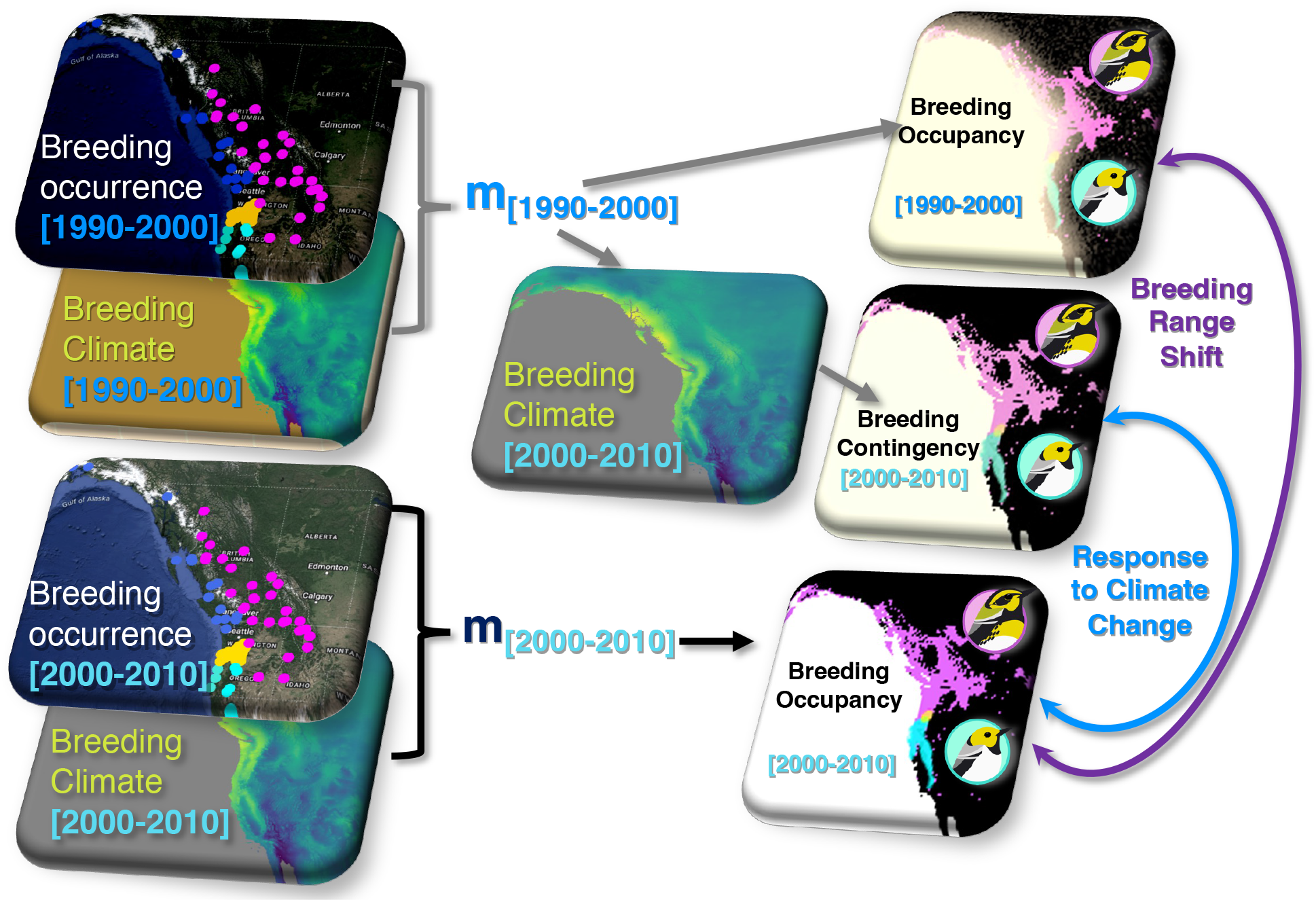
A schematic illustration of the tests of Breeding Range Shift and Organismal Response to Climate Change. **Organismal Response to Climate Change: Step 1**: calculate *expected breeding contingency* (*Exp*) with the historical model (e.g. m_[1990-2000]_) and the breeding climate of the current period (e.g. 2000-2010). **Step 2**: Calculate *predicted breeding occupancy* (*Pred*) with the recent model (e.g., m_[2000-2010]_). **Step 3**: compare *Pred* and *Exp*. A greater *Pred* than *Exp* reflects a positive response to climate change (e.g. climate adaptation). **Breeding Range Shift:** compare *Pred* within species across decades (e.g., 1990-2000 vs 2000-2010).

To test whether there was a significant difference between *expected breeding contingency* (*Exp*) and *predicted breeding occupancy* (*Pred*), I generated 1000 bootstrap samples each with 10,000 randomly sampled localities. For each bootstrap sample, I calculated *occupancy probability* as the number of localities with *breeding presence* (see above) over the total number of localities. The same set of localities applied for *Exp* and *Pred* extraction. With all the bootstrap samples, I ran a linear mixed effect model with the response variable being breeding occupancy probability, fixed effect being “Exp vs pred”, and random effect being bootstrap sample IDs. I compared the full model to the reduced model without the fixed effect to evaluate whether *Exp* and *Pred* were significantly different.

To further understand any potential knock-on effect on the breeding niche competition between SOCC and STOW, I compared the overlap of breeding occupancy across decades. All the analysis was conducted in R (R Core Team 2023). **Results**

For all three decades, the climate variables significantly predicted the breeding distribution of SOCC (R^2^ ≈0.41, accuracy ≈ 0.8, **Table 1**) and STOW (R^2^ ≈ 0.8, accuracy ≈ 0.96, **Table 1**).

**Table 1.**
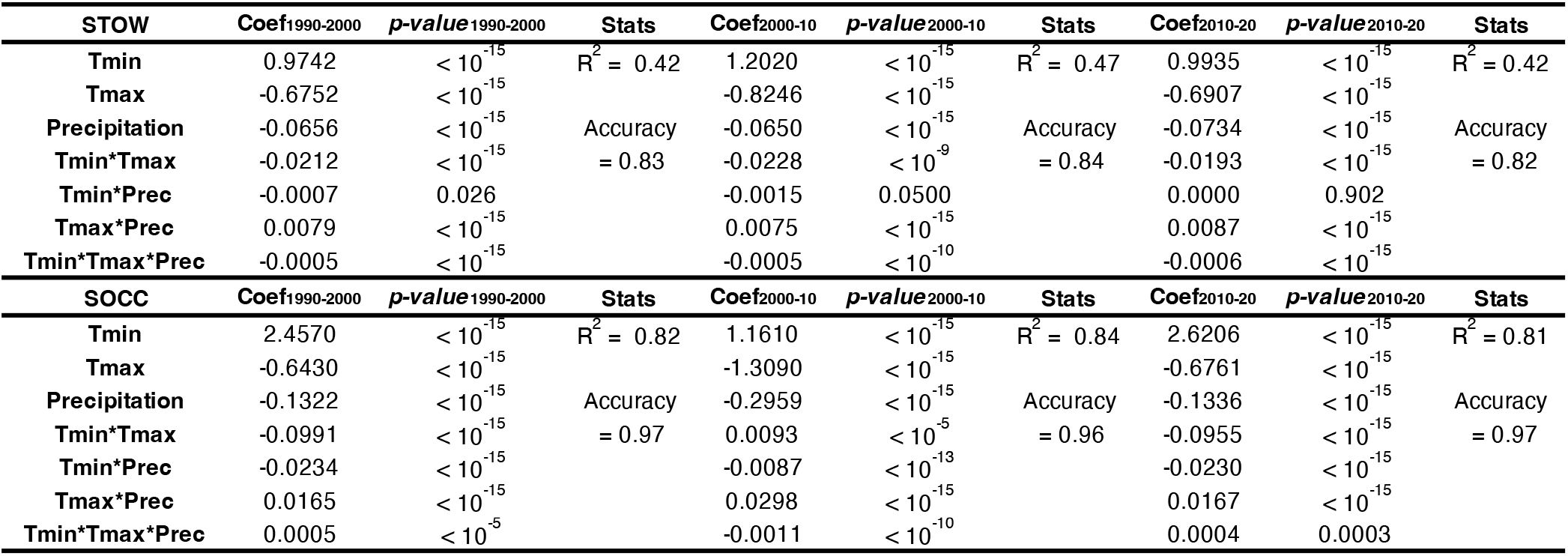
A summary of the breeding occupancy models across decades. Climate variables significantly explain breeding occupancy in STOW and SOCC in each decade m_[1990-2000],_ m_[2000-2010],_ m_[2010-2020]_.

I observed the lowest predicted breeding occupancy in 2000-10 for both SOCC and STOW (**Table 2**). The predicted breeding overlap between SOCC and STOW was also the lowest in 2000-10 (**Table 2, Fig. 5**). I detected the signature of breeding climate adaptation in 2010-20 for both species (**Fig. 2, 3; Table 3**) and STOW-only in 2000-10 (**Fig. 2; Table 3**). Specifically, in 2000-10, *Pred* was significantly greater than *Exp* in STOW (**Fig. 2 B, E, G**), but the reverse was true for SOCC (Table 3, **Fig. 3 B, E, G**). There was greater *Pred* than *Exp* in 2010-2020 for both STOW (**Table 3**; **Fig. 2 C, F, H**) and SOCC (**Table 3**; **Fig. 3 C, F, H**).

**Table 2.** The change in predicted breeding occupancy (*Pred*) of SOCC, STOW, and their breeding overlap in the past decades. Linear mixed effect model of the predicted presence among decades. The prediction was made with the occupancy model of each decade: m_[1990-2000],_ m_[2000-2010],_ m_[2010-2020]_.

| Species | Periods | Difference | 95%CI | p-value | Mixed Effect Model |
| --- | --- | --- | --- | --- | --- |
| SOCC | (1990-2000)-(2000-2010) | **0.00619** | (0.00611, 0.0063) | $p < 10^{-15}$ | |
| | (1990-2000)-(2010-2020) | **0.00318** | (0.00312, 0.00324) | $p < 10^{-15}$ | F = 174291 |
| | (2000-2010)-(2010-2020) | **-0.00301** | (-0.00306, -0.00295) | $p < 10^{-15}$ | $p < 10^{-15}$ |
| STOW | (1990-2000)-(2000-2010) | **0.01085** | (0.01073, 0.01098) | $p < 10^{-15}$ | |
| | (1990-2000)-(2010-2020) | **-0.00944** | (-0.00956, -0.00932) | $p < 10^{-15}$ | F = 496044 |
| | (2000-2010)-(2010-2020) | **-0.02029** | (-0.02042, -0.02016) | $p < 10^{-15}$ | $p < 10^{-15}$ |
| Overlap | (1990-2000)-(2000-2010) | **0.00082** | (0.00811, 0.00823) | $p < 10^{-15}$ | |
| | (1990-2000)-(2010-2020) | **-0.00180** | (-0.00187, -0.00179) | $p < 10^{-15}$ | F = 71545 |
| | (2000-2010)-(2010-2020) | **-0.01000** | (-0.01006, -0.00994) | $p < 10^{-15}$ | $p < 10^{-15}$ |

**Table 3.** The difference in the predicted probability between the expected contingency (*Exp*) and predicted occupancy (*Pred*) of STOW or SOCC in 2000-10 and 2010-20. Significant and negative mean differences between *Exp* and *Pred* indicate population potential of climate adaptation during the decade reflects a positive response to climate change. *Exp* calculated by the historical occupancy model (m_[1990-2000]_) and the climate variation in the respective decade (2000-10 or 2010-20). *Pred* is inferred by the occupancy model (m_[2000-10]_ or m_[2010-20])_ in the respective decade.

| Species | Probability | Mean | 95%CI | T<sub>9999</sub> | p-value |
| --- | --- | --- | --- | --- | --- |
| STOW | p2000-10 (exp - pred) | -0.0345 | (-0.0349, -0.0341) | -164.71 | < 2.2e-16 |
|  | p2010-20 (exp - pred) | -0.0038 | (-0.0042, -0.0034) | -18.47 | < 2.2e-16 |
| SOCC | p2000-10 (exp - pred) | 0.0104 | (0.0102, 0.0106) | 95.74 | < 2.2e-16 |
|  | p2010-20 (exp - pred) | -0.0065 | (-0.0067, -0.0063) | -52.97 | < 2.2e-16 |

**Fig. 2.**
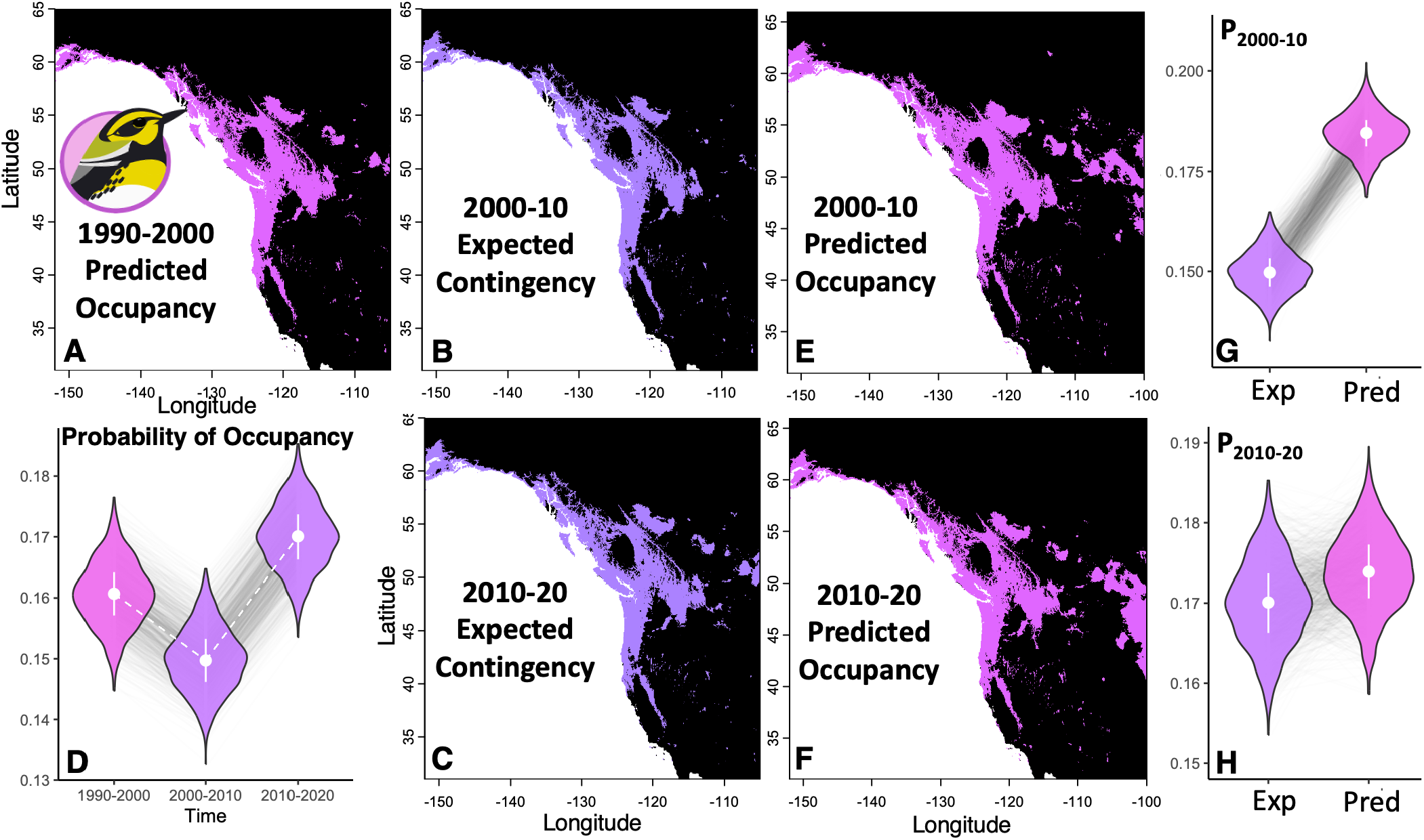
The greater predicted breeding occupancy (*Pred*) than expected continency (*Exp*) indicates a positive response to climate change in *Setophaga townsendi* (STOW). **B, E, G**: In 2000-10, the *Pred* (**E**) is greater than *Exp* (**B**). **C, F, H**: A similar pattern was observed in 2010-20, where the predicted breeding occupancy (**F**) is less than the expected contingency (**C**).

**Fig. 3.**
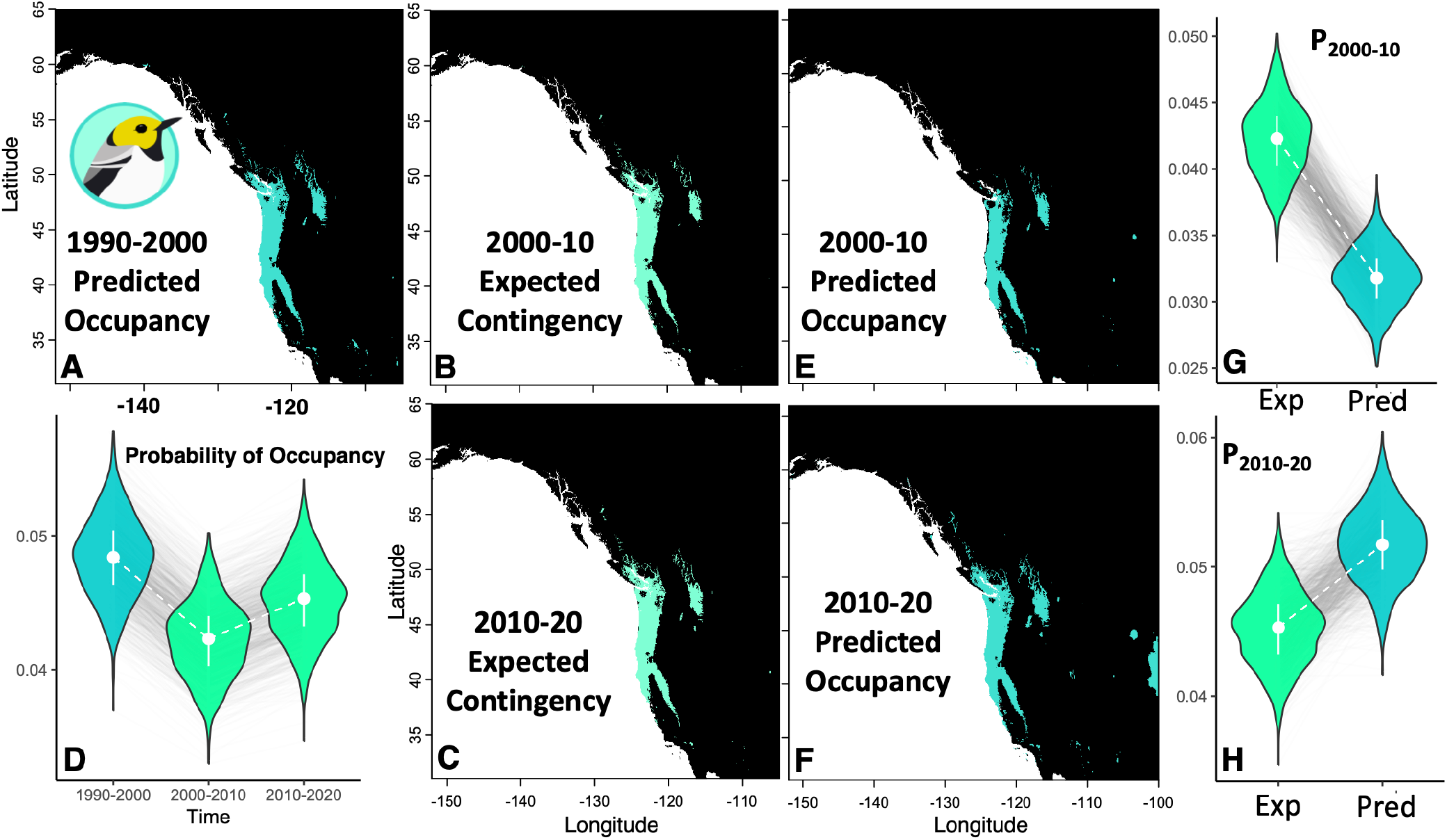
The heterogeneous response to climate change in *Setophaga occidentalis* (SOCC), as reflected by the difference between predicted breeding occupancy (*Pred*) and expected continency (*Exp*). **B, E, G**: In 2000-10, the *Pred* (**E**) is less than *Exp* (**B**). **C, F, H**: an opposite pattern was observed in 2010-20, where the *Pred* (**F**) was greater than *Exp* (**C**), which indicates climate adaptation in this decade.

## Discussion

Understanding the rainforest ecosystem response to climate change is central to global biodiversity conservation, as rainforests harbor the majority of terrestrial biodiversity (McMahon et al. 2011). An efficient and powerful predictive framework for dissecting rainforest indicator species’ response to climate change is essential. The rich citizen science data for the *Setophaga* warblers that breed in the temperate rainforest of North America empowered us to unravel the functional response of the rainforest to climate change. I observed a positive response to climate change in both species, as the recent breeding occupancy was greater than the expectation based on historical contingency (**Fig. 2, 3**). This pattern indicates breeding plasticity or adaptation to the changing climatic conditions in the breeding habitat. The decadal trend in the indicator species of the temperate rainforest could indicate the functional adaptability of the rainforest ecosystem to climate change.

The climate variables significantly explained the breeding distribution of both species (**Table 1**) with a strong effect on the breeding distribution of southern species (SOCC, R^2^ ∼ 0.8; **Fig. 3**) and a moderate effect on the northern species (STOW, R^2^ ∼ 0.4; **Fig. 2**). A caveat of our prediction framework is that it is based on limited climatic variables -- temperature and precipitation. Additional climate factors could elevate the predicting power. Despite that, the variation explained by two key climate factors was remarkable, especially for SOCC breeding occupancy.

More latent factors could contribute to STOW breeding occupancy, which spans over heterogeneous climatic conditions. The coastal STOW populations harbor mitonuclear ancestry from SOCC (Krosby and Rohwer 2009; Wang et al. 2021). The divergent SOCC and inland STOW mitonuclear ancestries have been associated with lipid metabolism in divergent climate adaptation (Wang et al. 2021). The predicted STOW breeding occupancy in the SOCC breeding range in southern Oregon and California (**Fig. 2**) could be attributed to the fact that coastal STOW contains mitonuclear variants of SOCC. The elevated genetic diversity from ancient hybridization in coastal British Columbia and Alaska (Wang et al. 2021) could predispose STOW with greater adaptive potential. Future studies could further dissect the heterogeneity in the breeding distribution of STOW concerning metabolic genotypes.

The excessive breeding occupancy than expected based on historical climatic reaction contingency could reflect climate adaptation. If so, the genetic variants suitable for the recent climatic conditions could have increased in frequency in the recent decade resulting in change in the niche adaptability. Alternatively, the observed positive response could be explained by breeding plasticity within the breeding niche potential. The historical climate-occupancy contingency may only represent the historical realized niche within a greater niche potential. However, the niche potential could also evolve and track climate change. Future studies could differentiate the possibilities by investigating any change in genetic frequency related to climate adaptation.

It is alarming that SOCC showed negative response to climate change in 2000-10 when the breeding occupancy was lower than expected based on niche contingency (**Fig. 3 G**). This pattern is consistent with their recent negative population trend of - 0.961 (95% CI: -1.675 to -0.285) (Sauer et al. 2020). Since annual precipitation is highly predictively of SOCC breeding occupancy (Table 1), the plummeting July precipitation since 2000-10 (**Fig. 4**) could be especially harmful. Despite that the July precipitation did not recover in 2010-20, SOCC showed signs of recovery, where the breeding occupancy surpassed expected breeding contingency. Further monitoring is need to determine whether this reversal could halt SOCC population decline.

**Fig. 4.**
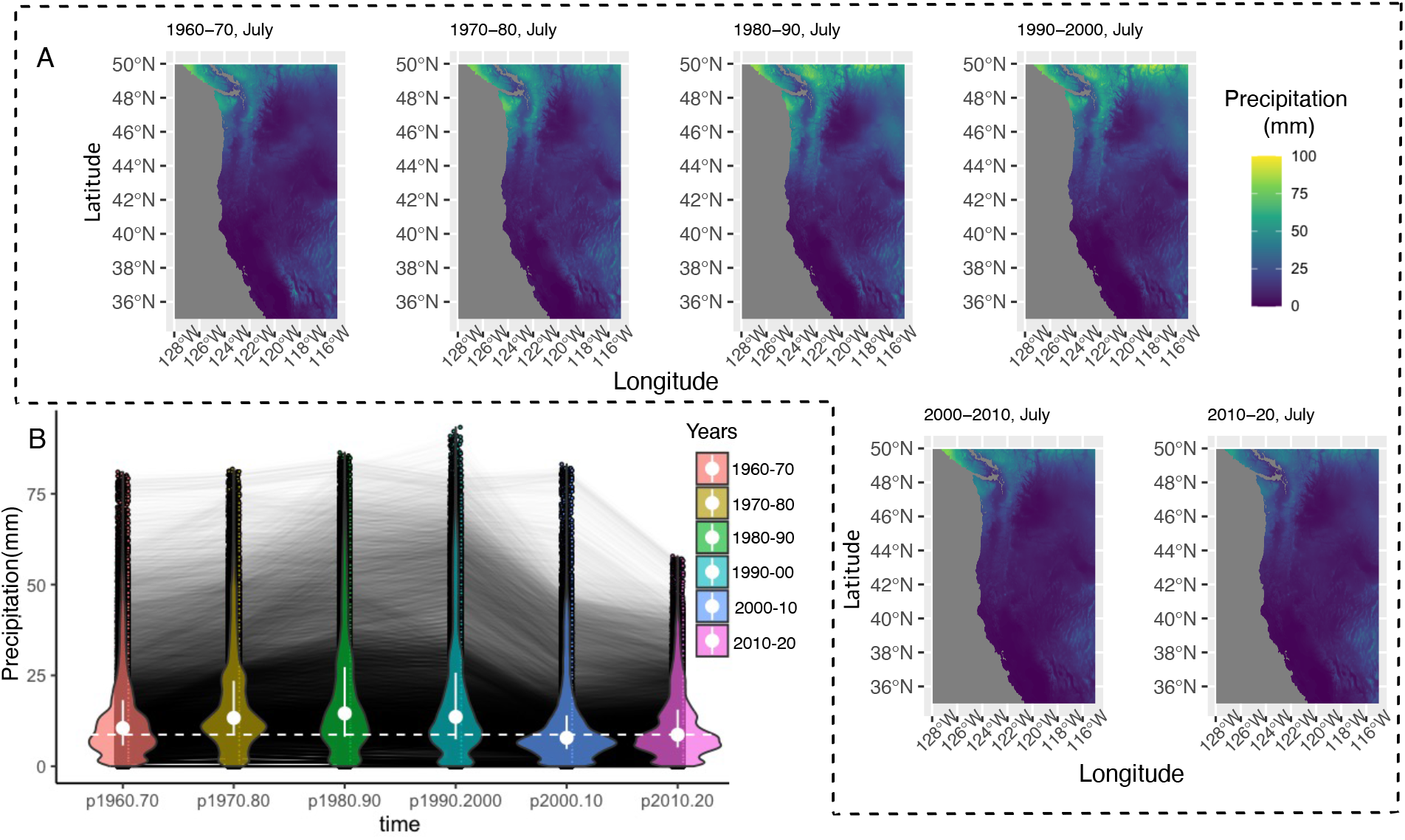
Change in mean July precipitation over decades. **A**, July mean precipitation (mm) over each decades from 1960-70 to 2010-20. **B**, violin plot showing the change of precipitation with each line being a location. There was a significant reduction of precipitation after 2000 (repeated measure ANOVA, post-hoc paired T-test with Bonferroni-corrected *p* < 0.05).

**Fig. 5.**
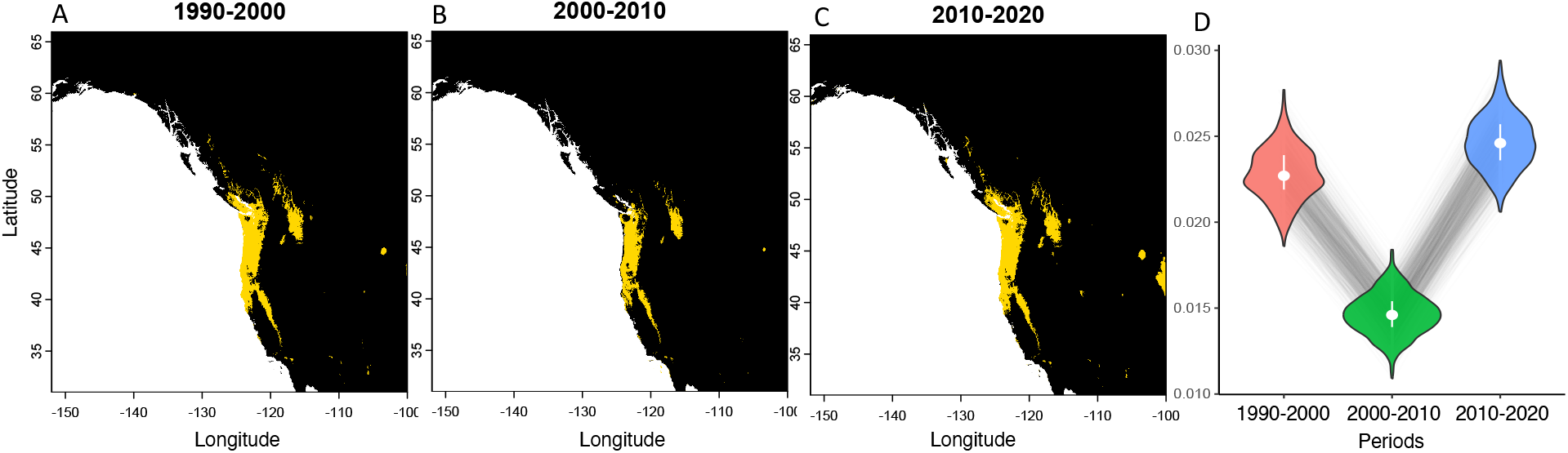
The change in the potential of breeding niche competition between SOCC and STOW reflected by the overlap of predicted breeding occupancy of over three decades. The lowest predicted overlap occurred in 2000-10, when July precipitation drastically declined.

The potential of breeding habitat competition between the hybridizing STOW and SOCC could be reflected by the overlap of their predicted breeding occupancy. The two warbler species are ecologically similar during breeding season as specialists in the canopy of coniferous forests (Morrison 1982). In the hybrid zones, interspecific competition does occur besides hybridization (Pearson 2000; Pearson and Manuwal 2000; de Zwaan et al. 2022). The predicted breeding overlap covers much of the breeding range of SOCC indicating that the absence of STOW breeding occupancy in California and Southern Oregon could be due to breeding niche competition with SOCC.

By leveraging citizen science data from three decades, I conducted a modeling synthesis for understanding the breeding wood warbler response to climate change which illuminates the functional reaction of the temperate rainforest of North America. Overall, the indicator bird species demonstrated positive responses to climate change, despite the challenges represented by the rampant decline of July precipitation since 2000. This result reflects the climate adaptation of the hermit warblers, rendering hope for the temperate rainforest ecosystem. However, more ecological factors in different trophic levels need to be further evaluated to fully understand the ecosystem response to climate change. Future investigation of the physiological, behavioral, and phenological mechanisms underlying the organismal response to climate change could advance our understanding of climate adaptation in rainforest biomes, where most terrestrial biodiversity resides.

